# Sex-specific relationship between melanopsin-dependent light sensitivity and chronotype across the lifespan

**DOI:** 10.1101/2025.10.17.681907

**Authors:** G van der Zwet, Z Bor, R Bos, R van Dorp, LM Pape, LCA van der Zwet, EHC van Dijk, H van de Stadt, EM McGlashan, SH Michel, L Kervezee

## Abstract

**Study objectives:** Light, acting primarily via melanopsin-mediated signaling, plays a central role in synchronising circadian rhythms. Individuals vary markedly in the sensitivity of their circadian system to light. Whether these differences contribute to the interindividual variability in chronotype, a behavioural manifestation of internal circadian timing, is unclear. The aim of this study was to determine the relationship between melanopsin-dependent light sensitivity and chronotype across the general population.

**Methods:** Participants (adults and children aged ≥ 8 years) were recruited in a science museum. Chronotype was determined using the µMCTQ and the post-illumination pupillary response (PIPR) to short and long-wavelength light stimuli was used as a measure of melanopsin-dependent light sensitivity. The relationship between PIPR and chronotype and their interaction with age and sex were assessed using multiple linear regression.

**Results:** Pupil recordings and questionnaires were available from 457 participants, including 284 adults and 173 children. In adults, the relationship between melanopsin-dependent light sensitivity and chronotype depends on sex and age: in young adult men, greater light sensitivity is linked to a significantly later chronotype, whereas it is significantly associated with an earlier chronotype in older adult women. In children, no evidence was found for a relationship between light sensitivity and chronotype.

**Conclusions:** These findings suggest that individual variation in light sensitivity interacts with sex and age-specific differences in the circadian system and light exposure behaviour to influence circadian timing. Light exposure recommendations should be personalised to take into account these sex and age-specific effects.

**Statement of significance:** While individuals differ widely in how their circadian system responds to light, to what extent these individual responses influence internal circadian timing remains unclear. By studying a large and diverse sample of children and adults, our findings reveal that the relationship between light sensitivity and chronotype, a behavioural manifestation of circadian timing, is shaped by both age and sex, offering a more comprehensive understanding of this relationship than previously recognised. Specifically, greater light sensitivity is linked to later chronotype in young men but to earlier chronotype in older women. These results reveal that the impact of light on circadian timing changes across the lifespan and will contribute to the development of personalised light exposure guidelines to promote health and well-being.

## Introduction

Circadian rhythms are endogenous, approximately 24-hour cycles in physiological and behavioural processes that enable organisms to anticipate and adapt to the daily light-dark cycle [1]. In humans, the timing of these rhythms varies considerably among individuals, which can manifest as differences in chronotype [2]. Chronotype reflects the timing of an individual’s daily rhythms including their internal circadian timing, ranging from early chronotypes (“early birds”) to late chronotypes (“night owls”)[3]. Chronotype is related to a number of lifestyle factors and health outcomes. For example, being an evening type is associated with increased alcohol intake and tobacco use[4], an increased risk of adverse metabolic outcomes, and depressive symptoms[5]. Therefore, it is critical to gain an understanding of the biological factors leading to the variation in chronotype across the population.

The factors contributing to interindividual differences in chronotype are multifaceted. Genetic factors play a role[2]: a genome-wide association study identified 351 genetic loci associated with self-reported morningness [6] and it was found that a missense mutation in one of the genes involved in the molecular clock machinery can lead to familial advanced sleep phase syndrome, manifested as an extreme early chronotype [7]. Chronotype changes across the lifespan [3], with typically early chronotypes in childhood, a progressively later chronotype over the course of adolescence, and a more gradual return to an earlier chronotype during adulthood [8–14]. Chronotype also depends on sex: men tend to have a later chronotype than women [15].

The variation in chronotype in the population may also be due to interindividual differences in the response to environmental timing cues that synchronize the circadian system to the external environment, such as light [16,17]. Individuals vastly differ in how sensitive their circadian system is to light, with a more than 50-fold difference in sensitivity across individuals [18]. Therefore, it is plausible that even with similar light exposure patterns, interindividual differences in sensitivity to light in the population may lead to differences in the synchronisation of the circadian system.

External light cues are conveyed to the central circadian clock in the suprachiasmatic nucleus (SCN) of the hypothalamus via intrinsically photosensitive retinal ganglion cells (ipRGCs) in the eye [19–21]. IpRGCs contain the photopigment melanopsin that is sensitive to short wavelengths of visible light (i.e., blue light) [22–24]. Besides synchronising the circadian system to the environment, melanopsin-mediated phototransduction via ipRGCs is known to mediate other non-visual effects of light[25], such as pupillary light responses via direct projections to the olivary pretectal nucleus [19]. Specifically, ipRGCs drive the post-illumination pupillary response (PIPR), the sustained constriction of the pupil after light offset, providing a non-invasive biomarker for assessing individual differences in melanopsin-mediated phototransduction.

It remains unclear whether and to which degree variation in melanopsin-mediated phototransduction can explain differences in chronotype across the general population. Addressing this question is essential to advance our understanding of the biological basis of chronotype variation [16]. In addition, given that light is the most dominant cue that synchronises the circadian system to the external environment and light exposure at the wrong time of day is associated with poor physical and mental health outcomes [26–28], insight into the factors that contribute to the interindividual variation in this relationship are essential to develop effective, personalised, light exposure recommendations.

Prior studies in specific cohorts provide some evidence for a positive association between circadian timing and PIPR. For example, in a cohort of young healthy adults, a larger PIPR was associated with later sleep timing [29], and in a cohort of adults with varying depression severity, a larger PIPR was associated with later circadian timing in summer but not in winter [30]. In this study, we investigate the relationship between PIPR and chronotype across a wide age range and in different sexes. Making use of the unique opportunity to conduct this study in a sample of science museum visitors, our goal was not only to elucidate the role of melanopsin-mediated phototransduction in shaping individual chronotype but also to promote awareness of circadian biology in a broad community, thereby bridging the gap between science and society.

## Methods

### Participants

This study was part of the NEMO Science Live program during a two-week school holiday. Participants were recruited and studied during museum opening hours (10:00-17:30) on an ongoing basis over 13 days during the Christmas holiday. Visitors of the NEMO Science Museum in Amsterdam, who were ≥ 8 years old and proficient in Dutch or English, were invited to participate in this study as part of their visit to the museum. Up to four participants could be accommodated per session allowing for groups of visitors (e.g. families) to participate together. Children younger than 18 years were accompanied by a parent or guardian to sign informed consent. All research data were collected anonymously. Participants were excluded from the study if they reported having epileptic disorders, using illicit drugs or cannabis in the past seven days, or using pupil-dilating eye drops in the past two weeks. In addition, they were excluded if they reported being unable or unwilling to be in a dimly lit room for ∼10 minutes (e.g., due to nyctophobia or claustrophobia), or if they reported experiencing discomfort after receiving a light stimulus in the dark (e.g., due to photophobia or migraine). The study protocol was approved by the Medical Ethics Committee (nWMO Commissie Divisie 4) of the Leiden University Medical Center under registration number nWMO-D4-2024-020.

### Study protocol

After providing written informed consent and passing the screening, participants were asked to fill out a questionnaire about basic demographics, general health and lifestyle (including smoking, caffeine, and alcohol intake), use of medication, sleep habits and ocular health. Chronotype was determined by the micro Munich Chronotype Questionnaire (μMCTQ).[31] For the questions in the μMCTQ related to sleep and wake timing on free days, participants were instructed to answer these questions with a day in mind that is free of commitments or disturbances, such as alarm clocks or house mates (e.g., young children), that may disrupt their sleep. A separate version of the questionnaire was used for children (participants below the age of 18), in which the questions about alcohol use, smoking, and night shift work were omitted and questions about chronotype were worded differently to be more applicable (e.g., school days instead of working days). Subsequently, participants were taken to a separate room to undergo the pupil recording. Prior to the start of the pupil recording, the participants were informed about each step of the experimental protocol, and the light stimuli were demonstrated to indicate the brightness to be expected. Subsequently, the main overhead lights were turned off and the participants underwent a dark adaptation period (< 2 photopic lux) of 5 minutes, in line with previous studies [32–35].

After dark adaptation, a monocular pupil recording of the right eye and the light exposure protocol were started. For the light protocol, baseline pupil diameter data were captured for 30 seconds in the dark, followed by a 1-second pulse of red light, 60 seconds of inter-stimulus darkness, a 1-second pulse of blue light, and 30 seconds of post-stimulus darkness (**Supplementary Figure S1A**). The red light stimulus served as a control to account for the contribution of other retinal photoreceptors to the PIPR. Participants were asked approximately 10 seconds prior to a light pulse to move and blink as little as possible until 10 seconds after light offset. After completion of the pupil recording, the room lights were turned on again and the participants were shown images of their pupil before and after the two light pulses (**Supplementary Figure S1B**).

### Light stimuli

Four identical experimental setups were developed that consisted of a custom-made rectangular light panel of 27 cm x 18 cm (width x height) that consisted of RGB LED array (NeoPixel WS2812, Adafruit, New York City, USA) covered by diffuser material with optimal light transmittance (PyraLed Makrolon Dx NR 139 cool, LT = 87%, Pyrasied, Leeuwarden, the Netherlands) (**Supplementary Figure S1C**). The light color and intensity of the light panel was controlled by an Adafruit Gemma controller (Adafruit, New York City, USA). Participant ID was entered on a laptop using a custom-made user interface (Python v 3.11.5). This Python interface was also used to control light exposures via an Arduino Uno microcontroller (Arduino, Monza, Italy) that triggered the RGB LEDs and to log the time stamps of light pulses, which was saved as a csv file after completion of the experimental protocol.

The pupil recording was performed in a dimly lit room (< 2 photopic lux). During the experiment, the distance between the participants’ cornea and the LED panel was 15 cm, yielding a visual angle of 84° x 62° (horizontal x vertical plane). The camera recorded the pupil through a cutout in the LED panel that covered approximately 20° of the central visual field. Irradiance at the level of the cornea was measured in the vertical plane for each of the experimental setups with a calibrated spectrometer (Avantes 1901283u1, Avantes, Apeldoorn, the Netherlands) using AvaSoftB software (AvaSoft 8.7.1.0 - 2017 Avantes) and converted to total irradiance and melanopic equivalent daylight illuminance (mEDI) using the CIE S 026 toolbox [36]. The red light had a total irradiance of 14.0 log photons/cm^2^/s, a mEDI of 1.3 lux, a photopic illuminance of 73 lux, and a peak wavelength of 627 nm with a bandwidth at half-max of 15 nm (**Supplementary Figure S1D**). The blue light had a total irradiance of 14.0 log photons/cm^2^/s, a mEDI of 244 lux, a photopic illuminance of 26 lux, and a peak wavelength of 463 nm with a bandwidth at half-max of 17 nm (**Supplementary Figure S1D**). Light conditions are reported in line with the ENLIGHT reporting guidelines [37].

### Pupil recordings

The pupil recordings were made using a custom-made setup, consisting of an infrared light source (M120, Kemo Electronic GmbH, Germany) and a monochrome camera (Basler daA1440-220um with Basler pylon 6.0.1 drivers) equipped with an infrared filter (IR850, 30.5mm, Green.L, China). Images were acquired using open-source PupilEXT software (beta version 0.1.1) [38]. Camera calibration was performed in PupilEXT with default settings. Throughout the experimental runs, the gain was set to 8 dB and the exposure time to 9 ms, while the aperture and focus of the camera were adapted to the individual participant to optimise brightness and contrast of the recording. The camera was set to capture images at 60 frames per second. For each image, the pupil diameter and the pupil outline confidence were determined by PupilEXT using the PuReST algorithm [39]. These data, along with the timestamps of the image acquisition, were stored in a csv file and used for furtheranalysis.

### Pupil data processing

All data processing and subsequent statistical analyses were performed in R v4.4.0 [40]. The light exposure protocol was matched to the pupil diameter time series using the recorded timestamps. Preprocessing of the raw pupil diameter time series was performed for each participant similar to the steps described by Martin *et al.* [41]. Firstly, to account for blinks, pupil diameter values were discarded if the first derivative of the diameter value exceeded ± 3 times the standard deviation or if the outline confidence was below 0.8. An additional preprocessing step was incorporated to account for cases in which the iris rather than the pupil was inadvertently detected by the algorithm, by setting values to missing if they were > 20% larger than the baseline value (i.e., the median pupil diameter 5 seconds before the red and blue lights were switched on). Missing values were replaced using linear interpolation. Timeseries were subsequently smoothened using a third-order Butterworth filter with a 4 Hz cutoff [41].

Next, time series were subjected to a quality control process to verify the reliability of the pupil recordings. Firstly, the stability of the timeseries was summarised by computing the relative median derivative of the smoothened timeseries at baseline and during the 6-second PIPR windows (between 5 and 7 seconds after lights off). If this value exceeded 20% (indicating instability of the recording, e.g., due to alternating detection of the pupil and the iris by the algorithm), the quality of the recording was deemed insufficient and the recording was excluded from further analysis. In addition, all pupil recordings were visually inspected independently by two authors (ZB and LK), along with the notes documented at the time of the experiment. Recordings were marked for exclusion if the experimental notes indicated a lack of compliance, accidental blinking during a light pulse, or if visual inspection revealed an unstable recording. Exclusions were then decided on by consensus between the two authors.

Following preprocessing, relative pupil diameter was calculated by expressing pupil diameter following the red and blue light onset as a percentage of the median (baseline) pupil diameter in the 5 seconds prior to the red and blue light onset, respectively. Subsequently, the net PIPR_6s_ was calculated from each timeseries by subtracting the median relative pupil diameter from 5 to 7 seconds after the blue light offset from the median relative pupil diameter from 5 to 7 seconds after the red light offset [42]. The PIPR_6s_ was used because it has least intra-individual variability compared to other PIPR metrics [43] and, despite showing circadian variation, it is minimally influenced by time of day during office hours (i.e., museum opening hours) [44–46].

### Statistical analysis

Data from adults (≥ 18 years old) and children (8-17 years old) were analyzed separately. As a first step, linear regression models were used to determine the effect of PIPR_6s_ (independent variable; continuous) on chronotype (dependent variable; continuous). Chronotype was determined using the µMCTQ by computing the midpoint of sleep on free days corrected for sleep debt (MSF_SC_) [31]. Chronotype is expressed in hours, with smaller values indicating early chronotypes (colloquially known as ‘morning types’) and larger values indicating late chronotypes (known as ‘evening types’). For all statistical analyses, main effects and interactions were considered significant if p < 0.05 (two-sided).

To assess if age and sex modulate the relationship between PIPR_6s_ and chronotype, linear regression models were used that included PIPR_6s_ (continuous variable), sex (categorical variable), and age (continuous variable) as main effects as well as interaction terms between PIPR_6s_ and age and between PIPR_6s_ and sex. Continuous variables were entered into the interaction models centered around their mean. Significance of model terms was assessed using Type II sum of squares analysis of variance tests using the R package car (v 3.1-3) [47]. Model assumptions (linearity and normality of residuals) were examined using histograms and Q-Q plots of the residuals.

To report the (back-transformed) model-estimated slopes and 95% confidence intervals (CIs) of the relationship between the chronotype and the independent variables and their interactions, the *emtrends* function from the R package emmeans (v1.11.1) [48] was used. The unit of the reported slopes and 95% CIs were expressed as the change in chronotype for each 10 percent point change in PIPR_6s_.To facilitate interpretation of the interaction effects, we determined the relationship between PIPR_6s_ and chronotype at three representative ages (for adults: 25, 40, and 55; for children: 9, 12, and 15), corresponding to the lower, middle, and upper ranges of the age distribution in the two populations. These ages were selected for interpretability and do not reflect categorisation of age in the statistical model. The slope at a certain sex category or age was considered significant if the 95% CI did not encompass 0.

A sensitivity analysis was carried out to examine to what extent the observed relationships are present in a ‘clean’ population that is more similar to study populations recruited for laboratory studies. To this end, the interaction model was fit to a dataset from which all participants were excluded who reported any of the following criteria: being color blind, working a night shift more than once in the past month, taking any sleep medication in the past night, having smoked or drunk alcohol on the day of the experiment, or drinking more than one caffeinated beverage on the day of the experiment. In addition, all participants that reported having ocular abnormalities or taking any medication in the past 24 hours that may affect the study outcomes (reviewed by study physician EvD) were excluded from this analysis. Similarly, a second sensitivity analysis was conducted to evaluate the impact of negative (i.e., potentially unphysiological) PIPR_6s_ values on the model estimates by excluding all PIPR_6s_ values smaller than 0 from the dataset. Model estimates for these sensitivity analyses were visualised as described above for the main analysis.

## Results

### Participants

Of the total of 612 participants that provided written informed consent and underwent the screening, data from 457 participants were available for analysis following data cleaning (see **Figure 1** for the inclusion flowchart), of which 284 were adults (≥ 18 years old) and 173 were children (8-17 years old). The total number of included participants per day ranged from 25 to 52, with a median of 35 recordings per day (**Supplementary Figure S2**). For detailed participant information, see **Table 1**. The age distribution of the entire study population was bimodal (**Figure 2A**), reflecting the expected demographic composition of science museum visitors during a school holiday, consisting predominantly of children and their parents. Children had an earlier chronotype (MSF_SC_: 2.86 [IQR: 2.16-3.50] h) than adults (3.46 [IQR: 2.96-4.14] h, t(366)=6.9, p < 0.001) (**Table 1**, **Figure 2B**). Of note, we found that chronotype depends on sex and age, with chronotype being generally earlier in women than in men and becoming progressively later from childhood through adolescence, then gradually shifting to earlier in adulthood (**Figure 2C**), visually similar to what has been observed previously in large epidemiological studies [8,14,49].

**Figure 1:**
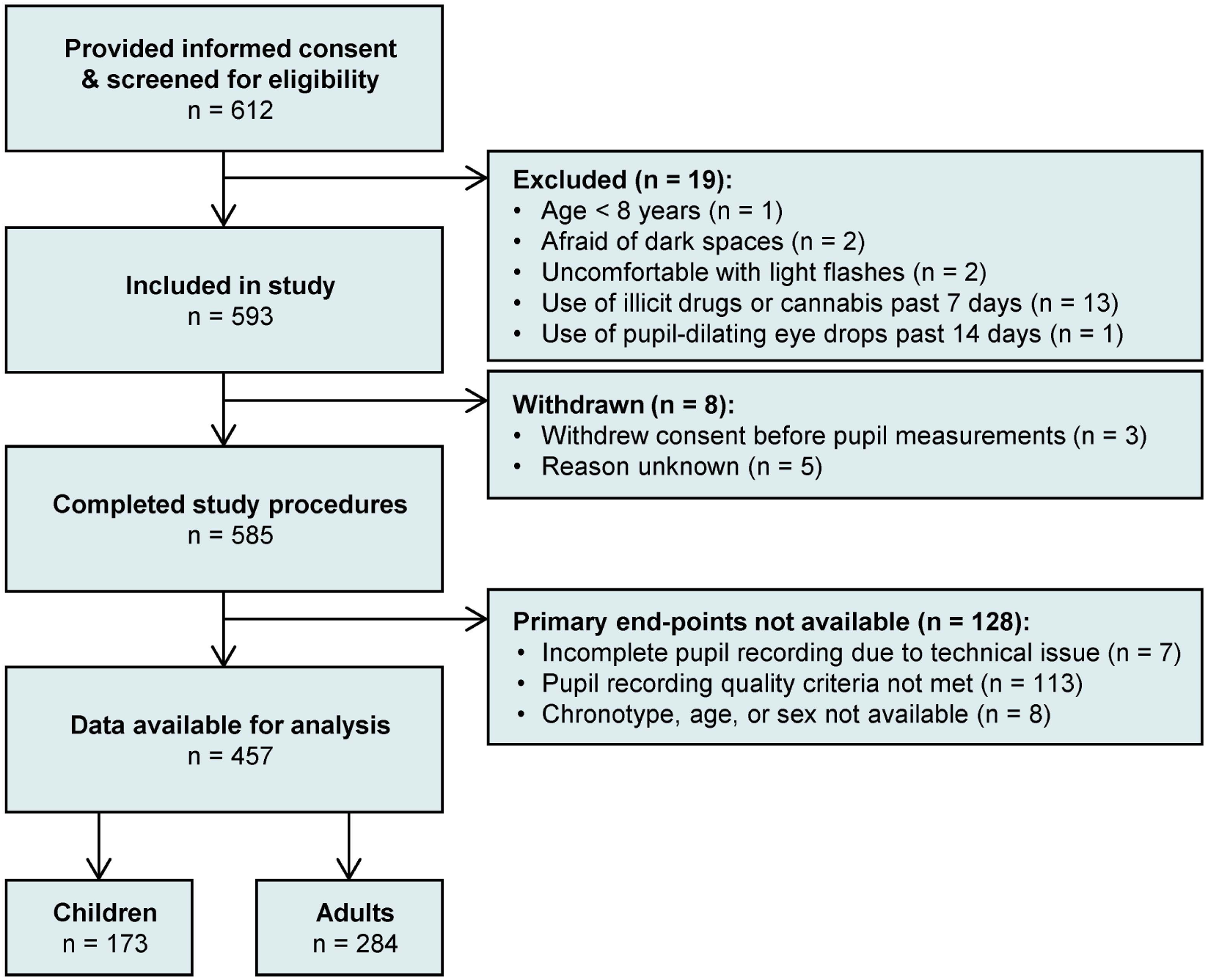
Flowchart of participant inclusion.

**Figure 2:**
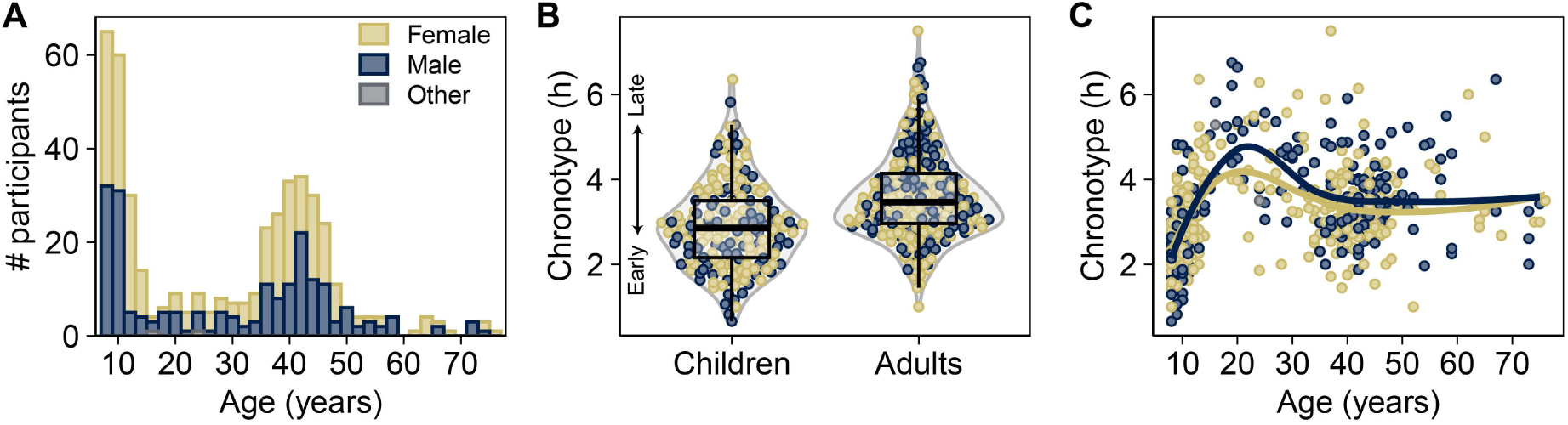
**Distribution of age and chronotype and their relationship**. **(A)** Age distribution of included participants, split in females (n = 250), males (n = 205) and others (n = 2). **(B)** Chronotype distribution in children (n = 173) and adults (n = 284). Boxplots are shown as median and interquartile ranges (IQR), with whiskers extending to 1.5 IQR. Chronotype is defined as the midpoint of sleep on free days corrected for sleep debt (MSF_SC_) in hours. **(C)** Relationship between age and chronotype (MSF_SC_) across sexes. Solid lines indicate smoothed curves fitted through the data for males and females separately using loess regression. Color coding in panel B and C as in panel A.

**Table 1:** Participant characteristics

| **Characteristic** | **Adults**<br>(n = 284) | **Children**<br>(n = 173) |
| --- | --- | --- |
| **Age (y)** | 41 (36, 46) | 10 (9, 12) |
| **Sex** |  |  |
| Female | 152 (54%) | 98 (57%) |
| Male | 131 (46%) | 74 (43%) |
| Other | 1 (0.4%) | 1 (0.6%) |
| **Chronotype (MSF<sub>sc</sub>; h)** | 3.46 (2.96, 4.14) | 2.86 (2.16, 3.50) |
| **Social jetlag (h)** | 0.75 (0.50, 1.38) | 1.00 (0.38, 1.67) |
| **Trouble sleeping prior night** | 60 (21%) | 41 (24%) |
| **Night shift work** | 18 (6.3%) | ND |
| **Smoking on experimental day** | 13 (4.6%)* | ND |
| **Alcohol consumption on experimental day** | 0 (0%)* | ND |
| **Caffeine intake on experimental day (number of cups)** |  |  |
| 0 | 64 (24%) | 153 (89%) |
| 1 | 89 (33%) | 17 (9.9%) |
| 2 | 67 (25%) | 2 (1.2%) |
| > 2 | 51 (19%) | 0 (0%) |
| Unknown | 13 | 1 |
| **Color blind** |  |  |
| No | 273 (97%) | 169 (98%) |
| Yes | 7 (2.5%) | 1 (0.6%) |
| Don't know | 2 (0.7%) | 2 (1.2%) |
| Unknown | 2 | 1 |

Data reported as median (Q1, Q3) or n (%). ND: not determined; MSFSC: midpoint of sleep on free days corrected for sleep debt.
\* Values missing from two participants

### Post-illumination pupillary response does not depend on time of day, age, or sex

Absolute baseline pupil diameters prior to the red and blue light stimuli were highly correlated (r^2^ = 0.95), with slightly lower baseline values before the blue light stimulus than before the red light stimulus (**Supplemental Figure S3**). As expected, pupil dilation following light offset was slower in response to the 1-second blue light stimulus than to the 1-second red light stimulus (**Figure 3A**), resulting in mostly positive values of the PIPR_6s_ (mean ± SEM: 10.3 ± 0.4%), with a large degree of interindividual variability (**Figure 3B**). No evidence was found for an effect of time of day on PIPR_6s_ (F(6, 450) = 0.699, p = 0.650, ANOVA; **Figure 3C**), nor of sex on PIPR_6s_ (t(423) = 0.65, p = 0.516, t-test, **Figure 3D**). In addition, no evidence was found that the PIPR_6s_ differed between adults and children (t(380) = -0.22, p = 0.827, t-test, Figure 3E).

**Figure 3:**
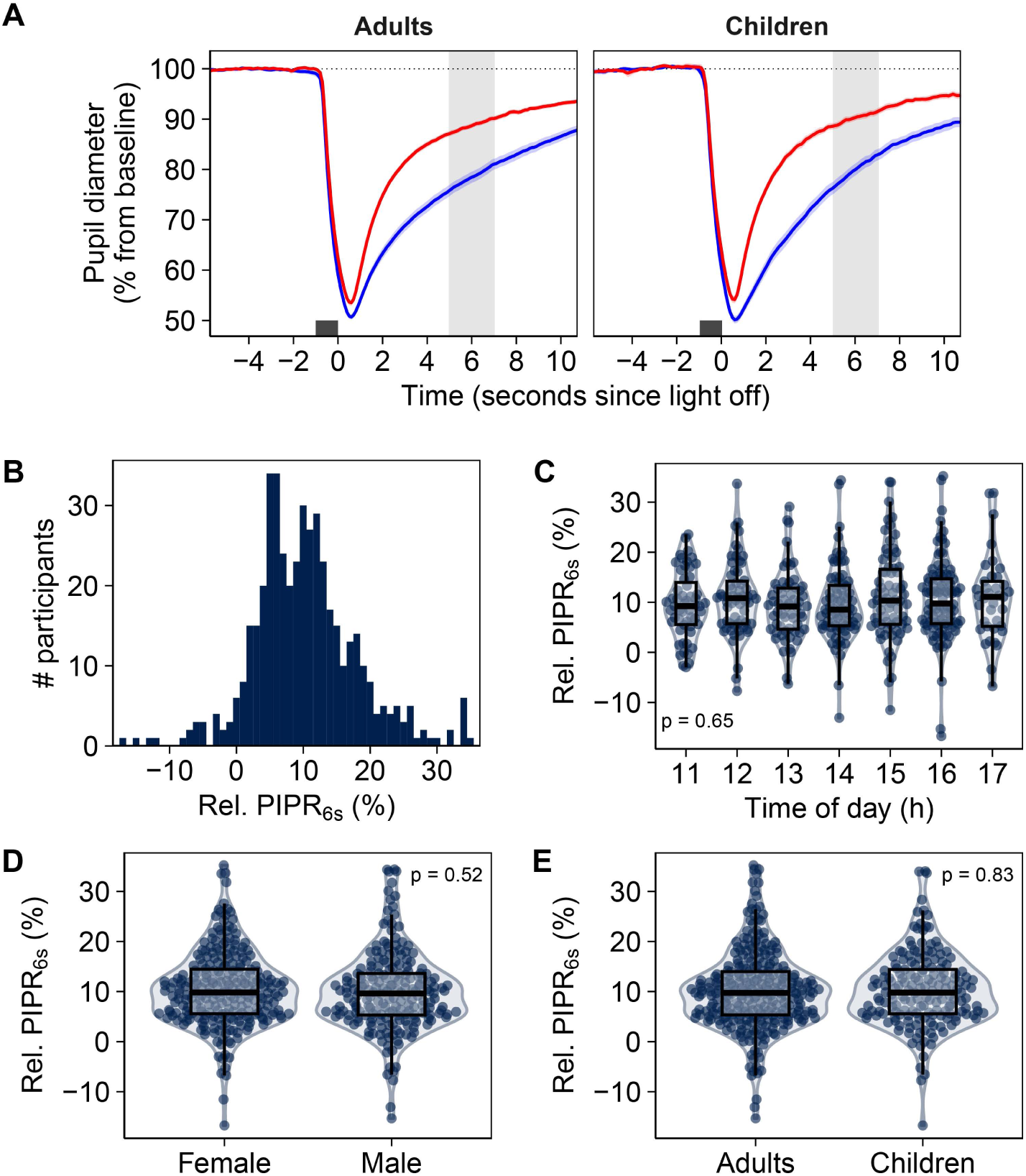
Effect of time of day, sex, and age on the post-illumination pupillary response. **(A**) Average pupil diameter time course in response to red and blue light in adults (n = 284) and children (n = 173). Data shown as mean and 95% CI. The dark grey box indicated the 1s light stimulus. Light grey box indicated the time window used to calculate the PIPR_6s_. **(B)** Distribution of relative PIPR_6s_ across all participants (n = 457). **(C)** PIPR_6s_ by time of day at which the pupil recording was performed across all participants (n = 457). Time of day was binned per hour (e.g., pupil recordings that started between 10:30 and 11:30 are categorised as 11). P-value indicates the significance of time of day on PIPR_6s_ based on one-way ANOVA. **(D)** PIPR_6s_ by sex (n = 455). Two participants who reported their sex as “other” were excluded from this panel. **(E)** PIPR_6s_ in adults and children (n = 457). P-values in panel C and D indicate the significance of sex and age on PIPR_6s_ based on student’s t-tests.

### Sex and age-specific relationship between chronotype and PIPR in adults

In adults, no significant overall relationship was found between chronotype and PIPR_6s_ (**Supplementary Figure S4A**). However, when including interaction effects to examine whether sex and age influence this relationship, we observed a significant interaction between PIPR_6s_ and sex on chronotype (p = 0.008) and a non-significant interaction between PIPR_6s_ and age on chronotype (p = 0.091) (**Table 2**). Post-hoc analyses to assess the relationship between PIPR_6s_ and chronotype in men and women at three representative ages revealed that, in women, a negative relationship between chronotype and PIPR_6s_ becomes more apparent with increasing age (**Figure 4A**). For the average woman at age 25, the relationship between PIPR_6s_ and chronotype is not significant (p = 0.532), while the association is significant at age 40 (p = 0.008) and age 55 (p = 0.002) (**Supplemental Table S1**). For women at age 40, the model predicts that for each 10 percent point increase in PIPR_6s_, chronotype is 17 minutes earlier. In men, a potentially positive relationship between chronotype and PIPR_6s_ is attenuated with increasing age (**Figure 4A**). At age 25, this relationship is significant (p = 0.036), whereas it is not significant at age 40 (p = 0.273) and age 55 (p = 0.670) (**Supplemental Table S1**).

**Figure 4:**
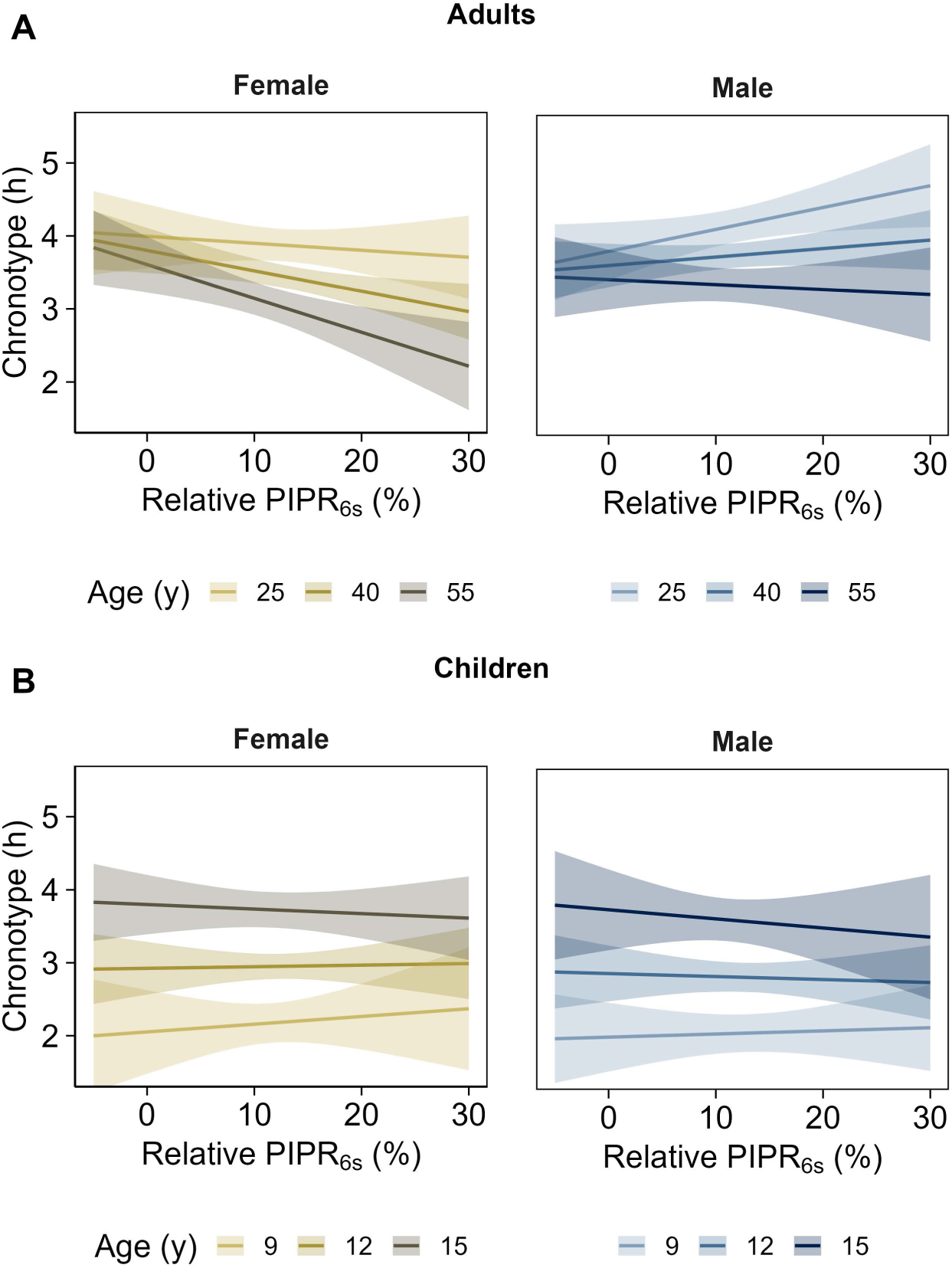
Model-estimated relationship between chronotype and post-illumination pupillary response (PIPR)_6s_ by sex at three representative ages in adults and children. **(A)** The model-estimated relationship between chronotype and PIPR_6s_ in adults shown for ages 25, 40, and 55 years, selected to represent the range of ages of the adult study population (18-76 years). **(B)** The model-estimated relationship between chronotype and PIPR_6s_ in children shown for ages 9, 12, and 15 years, selected to represent the range of ages of the child study population (8 – 18 years). Lines and shaded areas in panel A and B represent estimated marginal means and 95% confidence intervals at representative ages derived from the multiple linear regression model predicting chronotype by interactions between PIPR_6s_ and sex and PIPR_6s_ and age (modelled as a continuous variable).

**Table 2:** Effects of post-illumination pupillary response (PIPR), sex, age, and their interaction on chronotype in adults and children.

| Variables | Adults (n = 283)* |  | Children (n = 172)* |  |
| --- | --- | --- | --- | --- |
|  | F (df)<sup>#</sup> | P-value<sup>#</sup> | F (df)<sup>#</sup> | P-value<sup>#</sup> |
| PIPR<sub>6s</sub> | 1.08 (1, 277) | 0.300 | 0.02 (1, 166) | 0.880 |
| Sex | 8.05 (1, 277) | 0.005 | 0.64 (1, 166) | 0.426 |
| Age | 23.6 (1, 277) | <0.0001 | 66 (1, 166) | <0.0001 |
| PIPR<sub>6s</sub> x Sex | 7.14 (1, 277) | 0.008 | 0.10 (1, 166) | 0.754 |
| PIPR<sub>6s</sub> x Age | 2.88 (1, 277) | 0.091 | 0.41 (1, 166) | 0.524 |
|  | **Adjusted R<sup>2</sup>** |  | **Adjusted R<sup>2</sup>** |  |
| Explained variance | 0.114 |  | 0.280 |  |

\*Participants reporting their sex as 'Other' were excluded from these analysis (n = 1 in each age group); <sup>#</sup>F-statistics, degrees of freedom (df), and p-values obtained from type II sum-of-squares analysis of variance tests.

Since our study, performed in a science museum, was admittedly more noisy compared to the highly-controlled laboratory studies that typically examine the relationship between pupillary responses and the circadian system, a sensitivity analysis on data from a more homogenous subset of participants was performed. When excluding data from participants that meet exclusion criteria commonly applied in controlled laboratory studies (see Methods for the complete description), the direction and magnitude of the interaction effects between sex and PIPR_6s_ on chronotype is similar to that observed in the entire study population (**Supplementary Figure S5A**), supporting the robustness of our findings. Similarly, a sensitivity analysis excluding data from participants with a PIPR_6s_ value below 0 showed that our findings are minimally affected by these potentially unphysiological values (**Supplementary Figure S5B**).

### No relationship between chronotype and PIPR in children

In children, no overall significant relationship was found between chronotype and PIPR_6s_ (**Supplementary Figure S4B**). Furthermore, no influence of age or sex on this relationship was present: we observed no significant interaction between PIPR_6s_ and sex or PIPR_6s_ and age (**Table 2**). While the model-estimated relationship between chronotype and PIPR_6s_ shows no influence of sex and age on this relationship in children (**Figure 4B**), the main effect of age on chronotype is clearly visible, with the average 15-year old having a chronotype that is 1.5 [95% CI: 1.15 – 1.92] hours later than the average 9-year old (**Figure 4B**).

## Discussion

Individual differences in chronotype arise from a complex combination of biological and environmental factors, including sex, age, and genetics. However, the extent to which sensitivity of the circadian system to light, mediated by melanopsin-mediated phototransduction, contributes to these differences remains unclear. Making use of the unique opportunity to address this question among a large and diverse sample of science museum visitors, we reveal a sex-specific relationship between melanopsin-mediated light sensitivity and chronotype in adults, but not in children. Specifically, greater light sensitivity was associated with earlier chronotype in women, whereas this was not observed in men, highlighting the important role of sex in shaping the relationship between light sensitivity and chronotype in adults. Our results also show that age potentially modulates this relationship. As such, this study provides novel insights into how sex-specific differences in light sensitivity contribute to the variability of chronotype across the lifespan.

Without taking into account the effect of sex and age, no significant relationship between light sensitivity and chronotype was present in our data. We initially hypothesised that later chronotypes would have a larger PIPR, indicative of a circadian system that is more sensitive to light compared to earlier chronotypes. This hypothesis also fits with other lines of evidence suggesting that individual differences in the sensitivity of the circadian clock to light are related to the timing of their circadian system. For example, Wright *et al.* showed that in participants who were exposed to only natural light (daylight and campfire) for a week, the circadian system synchronises with the natural light-dark cycle [50]. This effect was particularly large in individuals with late circadian timing (as measured by the dim light melatonin onset), who shifted to an earlier circadian timing when exposed to natural light-dark regimes. These data suggest that the habitual exposure, and potentially greater sensitivity to, evening artificial light is what (partially) leads to their late circadian timing. Contrary to our initial hypothesis, we find that the relationship between light sensitivity and circadian timing is more complex and heavily depends on sex and possibly age. The finding that a larger PIPR is associated with an earlier chronotype in women but not in men could mechanistically be linked to previously identified sex differences in circadian function [15], such as differences in intrinsic circadian period [51] and light sensitivity of the circadian system [52]/ Duffy *et al.* found that the circadian period of women is six minutes shorter compared to men, which can account for differences in circadian timing between the sexes [51]. Furthermore, Vidafar et al. found that women exhibit a larger melatonin suppression than men under bright light (≥ 400 photopic lux) but not under moderate or dim light conditions (between 10 and 200 photopic lux) [52]. Consequently, the authors hypothesise that women might be more sensitive to the effects of bright (day)light on the circadian system than men. Together, a shorter circadian period and increased sensitivity to bright light suggest that women might be more sensitive to circadian phase advances than to phase delays, resulting in earlier circadian timing (e.g., chronotype) in more sensitive individuals, which is corroborated by our results. Future studies are necessary to validate the effect of sex differences on circadian phase shifting and its resulting consequences for chronotype variation between the sexes.

Our study also adds to the existing evidence that the non-visual effects of light depend on age [53]. We find that age, in addition to sex, may modulate the relationship between PIPR and chronotype in adults. Although this interaction effect was not significant in our main analysis and we had limited power to detect these effects due to the non-uniform distribution of age in our sample, the modulation by age warrants attention in future studies. Strikingly, when considering the combined effect of age and sex, we observe a more negative relationship between PIPR in women with increasing age, whereas young adult men have a more positive relationship between PIPR and chronotype which attenuates with increasing age. This effect of age may be (partly) explained by behavioural patterns of light exposure that differ across the lifespan [54]. Since our study took place in Amsterdam, we speculate that our adult study population consisted predominantly of parents of (young) children from near Amsterdam, where over 80% of school-aged children go to school on foot or by bike, often accompanied by their parents [55]. Thus, the many adults aged in their 40s in our sample are likely habitually exposed to substantial dose of high-intensity daylight in the morning. Mathematical models of the circadian system show that brighter daytime light conditions and decreased evening light exposure lead to earlier circadian phase and sleep timing [56–58]. Therefore, individuals with a higher light sensitivity who are exposed to light in the morning may have earlier circadian timing, and, consequently, an earlier chronotype. Notably, the accuracy of mathematical models of the circadian clock that consider behavioural light exposure is improved when making adjustments for both light sensitivity and intrinsic circadian period [58]. These results emphasise the highly nuanced interactions between underlying physiology (e.g., in circadian period or light sensitivity) and behavioural light exposure which should be considered in future investigations.

Although the magnitude of the PIPR itself did not differ between children and adults in our study, consistent with previous findings [59], the interaction with chronotype and sex that we observed in adults was not present in children. This discrepancy suggests developmental differences in the relationship between melanopsin-mediated phototransduction and chronotype, a topic that has received limited attention in children. Due to the enrichment for primary school-aged children and underrepresentation of pubescent and adolescent children in our sample, a potential differential effect of sex emerging during puberty might be absent in our data. Since sex differences that are amplified during puberty due to an increase in circulating sex hormones may influence circadian physiology [60], pre-pubertal children might exhibit a different relationship between PIPR and chronotype than adolescents and adults. Therefore, further research stratified by age, specifically in children and adolescents, is required to further examine the developmental changes in PIPR dynamics and light sensitivity of the circadian system during childhood and adolescence [61,62].

A strength of this study was its unique setting in a science museum, which allowed us to recruit a large and diverse sample in a relatively short time period. Importantly, the large number of participants willing to participate in our study during their museum visit emphasises the potential of using such a setting to promote awareness of circadian biology and the physiological effects of light in the wider community, while simultaneously collecting valuable scientific data. A limitation of this setting is that our study was relatively uncontrolled and designed to take up as little time as possible from the participants’ perspective so that it fitted within their museum visit. For example, the light stimuli were presented to the participants only once and were not repeated in case of blinks during light exposure or other artefacts. Although we tried to mitigate loss of data by demonstrating the light stimuli to the participants prior to dark adaptation and providing clear instructions, we had to exclude a relatively high number of pupil recordings due to insufficient quality (∼19%). In addition, another limitation is that the study population was substantially more heterogeneous than the controlled laboratory studies that are typically conducted in the field of chronobiology to study light sensitivity or circadian timing. While this heterogeneity increases the ecological validity of our findings, it may have introduced noise or confounders. However, the sensitivity analysis on a more homogenous subset of participants showed the same direction of the sex and age effects, supporting the robustness of our findings. Furthermore, our study was conducted near the winter solstice, during which the availability of daylight in Amsterdam is limited (approximately 7 hours and 45 minutes). Since season was found to be an important determinant of the relationship between PIPR and circadian timing [30], it remains to be investigated to what extent our findings apply in other seasons. Given that our observed interactions may be partially driven by differences in habitual light exposure, it will be particularly interesting to investigate the degree to which they persist or change under changing day lengths across the year.

In summary, we show that sex and age influence the relationship between melanopsin-mediated phototransduction and chronotype, providing novel evidence on the highly dynamic interplay between sensitivity of the circadian clock to light and chronotype. In women, a greater light sensitivity is associated with an earlier chronotype, while this was not observed in men and children. This suggests that individual variation in light sensitivity interacts with other factors such as sex-specific differences in circadian rhythms or behaviour, to influence circadian timing. The striking observation that greater sensitivity to light may have the opposite relationship with chronotype in young adult men compared to older adult women calls for further research into the effect of age, and age-related changes in behaviour that influence daily light exposure patterns, on this relationship. Altogether, our study contributes to our understanding of how age and sex shape the relationship between light sensitivity and the circadian system and supports the development of individualised light exposure recommendations.

## Acknowledgments

The authors thank Marit van Gent for her assistance with digitalising the survey responses, and the NEMO science museum and its visitors for their participation.

---

**Supplementary Figure S1.**
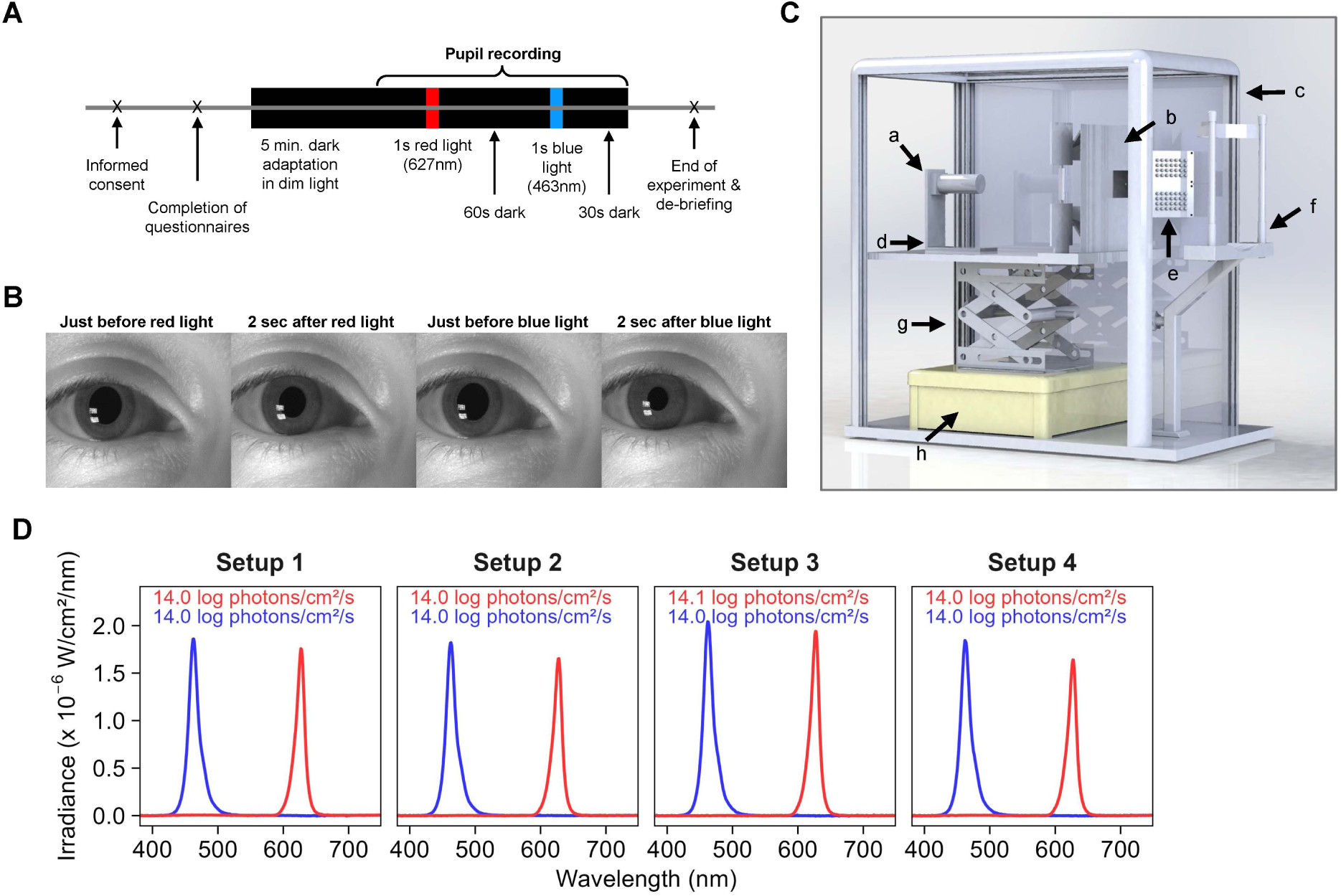
Study design. **(A)** Overview of the experimental protocol. **(B)** Example of the personalized images shown to participants as feedback at the end of the experimental session. **(C)** Three-dimensional rendering of the experimental setup with labels indicating the different components. a: monochrome camera; b: light panel (RGB LEDs covered with diffuser material); c: casing around the setup; d: horizontal adjustable platform; e: infrared light source; f: headrest; g: vertical adjustable platform; h: casing containing controller. **(D)** Spectral irradiance distribution of the red and blue light stimuli generated by each of the four experimental setups. Text insets show total irradiance per setup for the red (top) and blue (bottom) light stimuli.

**Supplemental Figure S2:**
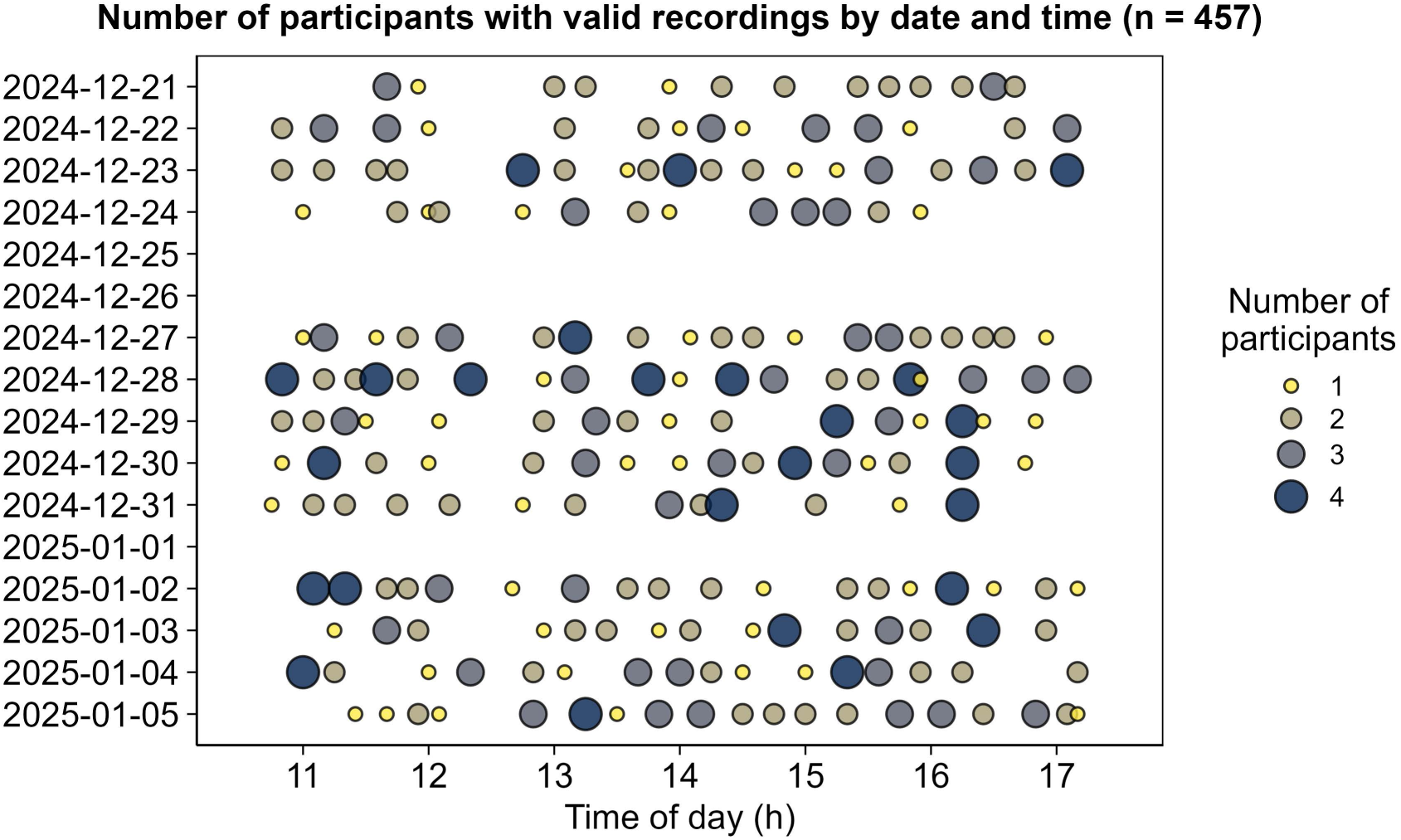
Number of participants by date and time. The experimental setup allowed for measuring up to four individuals simultaneously, resulting in the measurement of one to four participants per session.

**Supplemental Figure S3:**
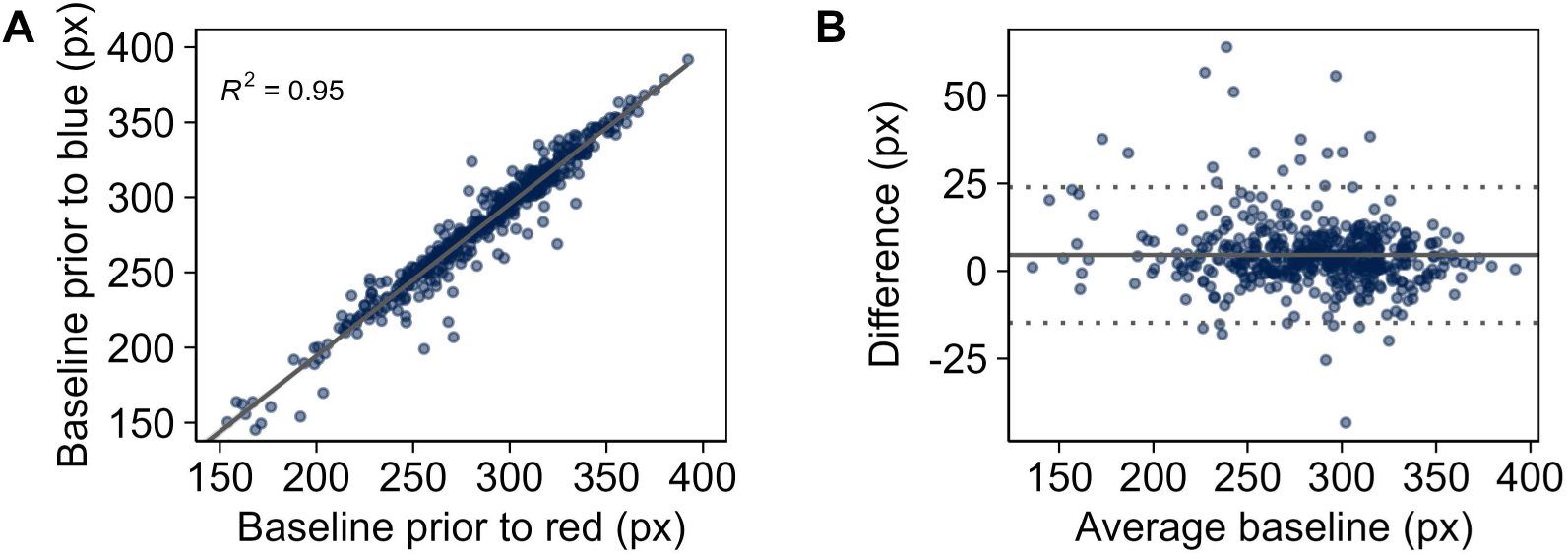
Comparison of baseline pupil diameter prior to the red and blue light stimuli. **(A)** Relationship between baseline pupil diameter before the red light pulse and the baseline pupil diameter before the blue light pulse in all participants (n = 457). Pupil diameters are shown in pixels. The solid line represents the linear regression line through the data points. The corresponding R^2^ value is shown in the plot. **(B)** Bland-Altman plot of the average of the baseline pupil diameter (in pixels) prior to the red and blue light pulses against the difference between the baseline pupil diameters before the red and the blue light pulses in all participants. The solid line represents the average difference in baseline, the dashed lines indicate 95% limits of agreement.

**Supplemental Figure S4:**
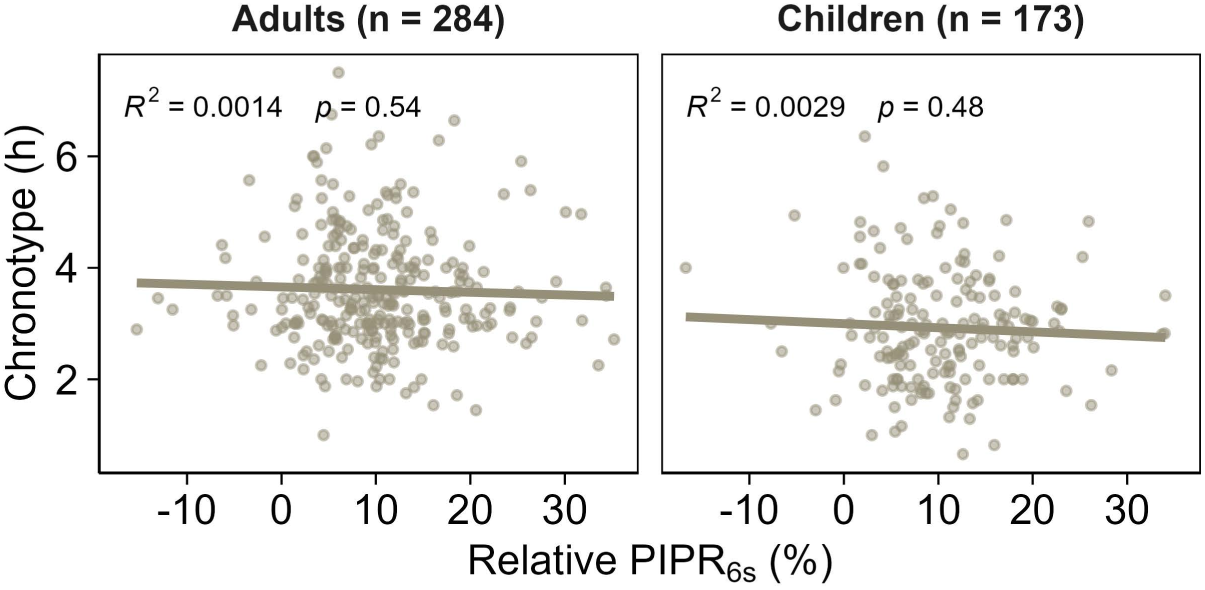
The correlation between chronotype and the relative post-illumination pupillary response (PIPR) at 6 seconds after the light offset in the raw data. The analysis is separated for children and adults. The dots represent the individuals. The line represents the correlation between chronotype and relative PIPR_6s_. The corresponding p values and R^2^ values are shown in the plots. This figure does not account for the effect of sex and age.

**Supplemental Figure S5.**
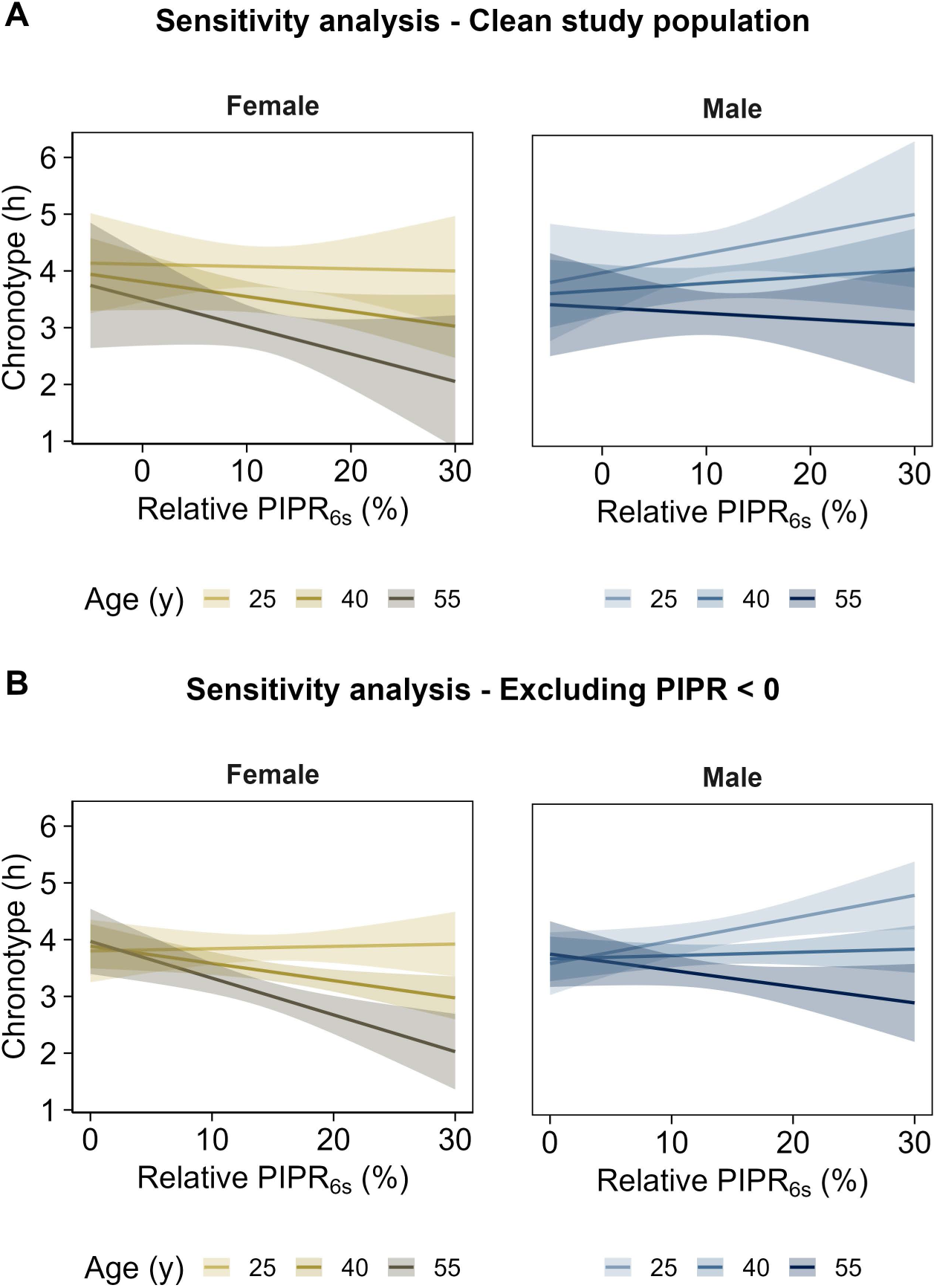
: Sensitivity analyses. **(A)** Model-estimated relationships between the post-illumination pupillary response (PIPR)_6s_ and chronotype at three representative ages in adult males and females in a ‘clean’ study population (see Methods for a full description of this subset). In total, 123 participants (out of the 283 adult participants) were included in this analysis (71 women, 52 men). Although the interaction effects are no longer significant (PIPR_6s_ x Sex: F(1, 117) = 2.54, p = 0.114; PIPR_6s_ x Age: F(1, 117) = 0.921, p = 0.339), the direction and magnitude of the effects are similar to that of the main analysis (see Figure 4 in the main text). **(B)** Model-estimated relationships between PIPR_6s_ and chronotype at three representative ages in adult males and females in the dataset excluding participants with a negative PIPR_6s_ value. In total, 269 participants were included in this analysis (145 women, 124 men). In this analysis, both interaction effects are significant (PIPR_6s_ x Sex: F(1, 263) = 4.53, p = 0.034; PIPR_6s_ x Age: F(1, 263) = 5.92, p = 0.016) and the direction and magnitude of the effects are similar to that of the main analysis.

**Supplemental Table S1:**
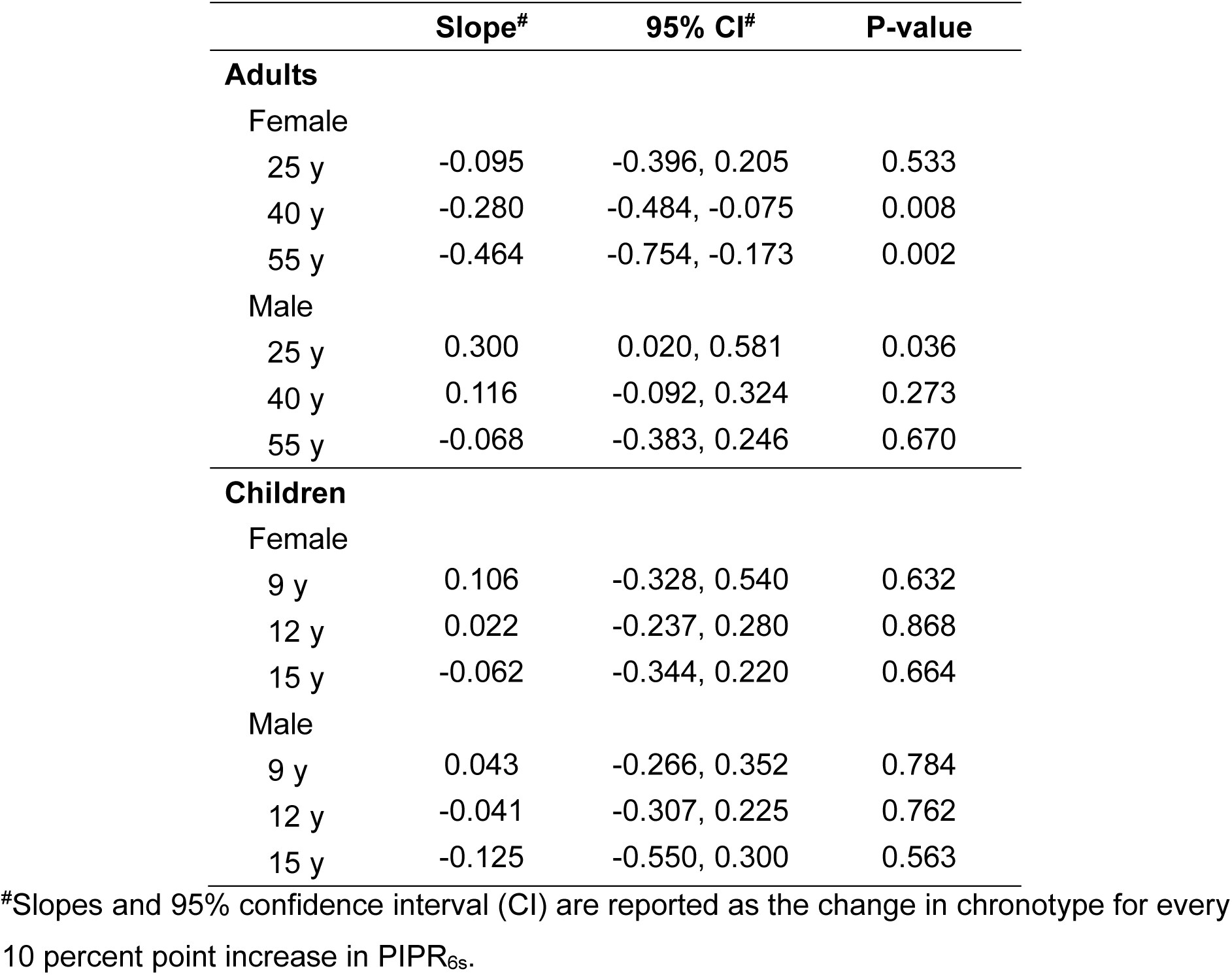
Estimated marginal means of the model-predicted slopes that describe the relationship between the post-illumination pupillary response (PIPR)_6s_ and chronotype by sex at three representative ages .

|  | **Slope<sup>#</sup>** | **95% CI<sup>#</sup>** | **P-value** |
| --- | --- | --- | --- |
| **Adults** |  |  |  |
| Female |  |  |  |
| 25 y | -0.095 | -0.396, 0.205 | 0.533 |
| 40 y | -0.280 | -0.484, -0.075 | 0.008 |
| 55 y | -0.464 | -0.754, -0.173 | 0.002 |
| Male |  |  |  |
| 25 y | 0.300 | 0.020, 0.581 | 0.036 |
| 40 y | 0.116 | -0.092, 0.324 | 0.273 |
| 55 y | -0.068 | -0.383, 0.246 | 0.670 |
| **Children** |  |  |  |
| Female |  |  |  |
| 9 y | 0.106 | -0.328, 0.540 | 0.632 |
| 12 y | 0.022 | -0.237, 0.280 | 0.868 |
| 15 y | -0.062 | -0.344, 0.220 | 0.664 |
| Male |  |  |  |
| 9 y | 0.043 | -0.266, 0.352 | 0.784 |
| 12 y | -0.041 | -0.307, 0.225 | 0.762 |
| 15 y | -0.125 | -0.550, 0.300 | 0.563 |

<sup>#</sup>Slopes and 95% confidence interval (CI) are reported as the change in chronotype for every 10 percent point increase in PIPR<sub>6s</sub>.

